# Identification of genetic factors controlling domestication-related traits in cowpea (*Vigna unguiculata* L. Walp)

**DOI:** 10.1101/202044

**Authors:** Sassoum Lo, María Muñoz-Amatriaín, Ousmane Boukar, Ira Herniter, Ndiaga Cisse, Yi-Ning Guo, Philip A. Roberts, Shizhong Xu, Christian Fatokun, Timothy J. Close

## Abstract

Cowpea (*Vigna unguiculata* L. Walp) is a warm-season legume with a genetically diverse gene-pool composed of wild and cultivated forms. Cowpea domestication involved considerable phenotypic changes from the wild progenitor, including reduction of pod shattering, increased organ size, and changes in flowering time. Little is known about the genetic basis underlying these changes. In this study, 215 recombinant inbred lines derived from a cross between a cultivated and a wild cowpea accession were used to evaluate nine domestication-related traits (pod shattering, peduncle length, flower color, flowering time, 100-seed weight, pod length, leaf length, leaf width and seed number per pod). A high-density genetic map containing 17,739 single nucleotide polymorphisms was constructed and used to identify 16 quantitative trait loci (QTL) for these nine domestication-related traits. Candidate genes underlying each of those 16 QTL were identified. Four regions with clusters of QTL were identified, including one on chromosome 8 related to increased organ size. This study provides new knowledge of the genomic regions controlling domestication-related traits in cowpea as well as candidate genes underlying those QTL. This information can help to exploit wild relatives in cowpea breeding programs.

**Key message:** This study identified regions of the cowpea genome that played an important role in cowpea domestication, including a hotspot region for increased organ size

## Introduction

Plant domestication is considered to be the key technological element of a transition to agriculture (Gepts 2014). Domesticated crop plants differ from their wild progenitors in various characters collectively termed the “domestication syndrome” (Hammer 1984). Some of the most important syndrome traits include loss of seed dispersal, early flowering, increased size of the edible organs, and changes in plant architecture and organ coloration (Abbo et al. 2014), and are the outcome of selection for adaptation to cultivated environments (Gepts 2010). The genetic control of domestication-related traits (DRTs) has been investigated in numerous crop species, including legumes, mainly by quantitative trait loci (QTL) mapping, using crosses between cultivated and wild accessions (Doebley et al. 1995; Frary et al. 2000; Isemura et al. 2007; Kaga et al. 2008; Koinange et al. 1996; Liu et al. 2007). Increased knowledge of the genomic regions controlling DRTs is valuable to efficiently exploit wild germplasm for improving biotic and abiotic stresses and overall yield.

Cowpea (*Vigna unguiculata* L. Walp) is one of the main sources of dietary protein and folic acid for millions of people in sub-Saharan Africa. Cowpea is a diploid (2n= 22) with a genetically diverse gene-pool (Pasquet 1999). The wild genepool is composed of both perennial and annual types, while cultivated forms are all annual. There are several cultivar groups, from which the most important are subsp. *unguiculata* (grain-type cowpea) and subsp. *sesquipedalis* (yardlong bean) (Marechal et al. 1978). *V. unguiculata* subsp. *dekindtiana,* a perennial form that only exists in Africa, is considered the wild progenitor of cowpea (Lush and Evans 1981). However, there is a lack of consensus on where in Africa cowpea domestication occurred. Several domestication locales have been proposed including West and Southeast Africa (Ba et al. 2004; Faris 1965; Rawal 1975; Steele and Mehra 1980; Vaillancourt and Weeden 1992).

Domestication of cowpea has, in general, resulted in a determinate growth habit, increased pod and seed size, early flowering, and reduction of pod shattering. Cultivated cowpea also shows a wide range of flower and seed coat colors, whereas wild cowpeas all have purple flowers and black seed coats. Only a few studies have identified QTL controlling DRTs in cowpea. Andargie et al. (2014) reported QTL for six DRTs including pod shattering, days to flowering, ovule number and growth habit. In subsp. *sesquipedalis*, QTL for DRTs including pod shattering, pod length and flower color have been also reported (Kongjaimun et al. 2012; Suanum et al. 2016; Xu et al. 2011; Xu et al. 2017). However, to date no candidate genes underlying domestication traits have been reported in cowpea.

In the present study, we investigated the genetic control of nine DRTs in cowpea including pod shattering, seed size, and flowering time using a population of 215 recombinant inbred lines (RILs) derived from a cross between a cultivated and a wild cowpea. The Cowpea iSelect Consortium Array, including 51,128 single nucleotide polymorphism (SNPs) markers (Muñoz-Amatriaín et al. 2017), was used to genotype this population, which allowed for high density genetic mapping. This study provides new insights into the genetic control of domestication-related traits in cowpea and can help to utilize more efficiently wild germplasm in breeding programs.

## Materials and Methods

### Plant material and growth conditions

A biparental mapping population of 215 F8 RILs derived from a cross between IT99K-573-1-1 and TVNu-1158, cultivated and wild cowpea accessions, respectively, was used for linkage mapping and QTL analysis. IT99K-573-1-1 is an early-maturing, white-seeded, high yielding and Striga resistant variety that was released in Nigeria under the name SAMPEA 14. TVNu-1158 is small seeded and has a perennial growth habit. It is cross-compatible with cultivated cowpea, although the F1 and subsequent generations showed partial sterility.

The parents and the F8 population were grown in pots filled with 5.0 kg topsoil and placed in a screen house at the International Institute of Tropical Agriculture (IITA), Ibadan, Nigeria (latitude 7° 30’N and longitude 3° 54’E, elevation 240 masl). Five seeds of each RIL were sown per pot and then thinned to two plant per pot when they were well established. The pots were watered regularly.

### Phenotypic data collection and analyses

A total of nine DRTs segregating in the population were evaluated (Table 1). Phenotypic data for pod shattering were collected by visual scoring, with scores of 0 = “no shattering”, and 1 = “pods opened and twisted”. Peduncle length was determined by measuring the distance from the point of peduncle attachment to the node on the stem to where the first flower bud emerged. Five peduncles were measured on each plant beginning with the lowest peduncle. Flower color was evaluated by visual scoring with 0 = “white”, and 1 = “purple”. Flowering time was scored as the number of days from date of planting to when the first flower opened. Seed weight was determined by the weight in grams of 100 seeds. The length (from base to the tip) and width (at the widest point) of the first leaf (opposite in arrangement) following seedling emergence was determined using a measuring tape. Pod length was determined by measuring the length of the first five pods per plant. Seed number per pod was evaluated by counting the number of seeds per 10 pods for each plant.

**Table 1:** Domestication-related traits evaluated in the F8 RILs population.

| General attribute | Organ | Trait |
| --- | --- | --- |
| Seed dispersal | Pod | Pod shattering |
| Plant type | Peduncle | Peduncle length |
| Pigmentation | Flower | Flower color |
| Phenology | Flower | Flowering time |
| Gigantism | Seed | 100-seed weight |
|  | Pod | Pod length |
|  | Leaf | Leaf length |
|  | Leaf | Leaf width |
| Yield potential | Seed | N° seeds per pod |

For the seven non binary traits, the mean, standard error, and range were calculated as well as the degrees of skewness. The frequency distribution of the seven traits was determined. The segregation pattern for pod shattering and flower color was analyzed by chi-square test for goodness-of-fit.

### Genotyping

Young leaves from each RIL and the parents were collected and placed into a ziplock bag containing silica gel packs for desiccation. Total genomic DNA of each line and parents was extracted from dried leaves using Plant DNeasy (Qiagen, Germany), quantified using Quant-IT dsDNA Assay Kit (Thermo Fisher Scientific, USA), and the concentration adjusted to 80 ng/μl. Genotyping was performed at the University of Southern California (Los Angeles, CA, USA) using the Cowpea iSelect Consortium Array, which includes 51,128 SNPs (Muñoz-Amatriaín et al. 2017).

### Linkage map construction

SNPs were called using the GenomeStudio software V.2011.1 (Illumina, Inc., San Diego, CA, USA). Data curation was performed by removing SNPs with more than 20% missing or heterozygous calls. Individuals were then plotted by missing calls and by heterozygous calls, and outliers were eliminated from further analysis. Lines carrying non-parental alleles and duplicated lines were also removed. Only SNPs that were polymorphic in both the parents and the RIL population, and had minor allele frequencies (MAFs) > 0.25 were used for linkage mapping. MSTmap (Wu et al. 2008; http://www.mstmap.org/) was used for genetic map construction, with the following parameters: grouping LOD criteria = 10; population type = DH (doubled haploid); no mapping size threshold = 2; no mapping distance threshold: 10 cM; try to detect genotyping errors = no; and genetic mapping function = kosambi. The chromosomes were numbered and oriented according to the recently developed cowpea pseudomolecules (Lonardi et al. 2017; https://phytozome.jgi.doe.gov/pz/portal.html#!info?alias=OrgVunguiculataer). Since the use of DH function inflated the cM distance for a RIL population, cM map distances were divided by two to correct for the extra round of effective recombination occurring in a RIL population compared to a DH population.

### QTL analysis and identification of candidate genes

QTL analysis was performed using a linear mixed model described by Xu (2013). In this approach, the background effect is captured via a polygene and a marker inferred kinship matrix (Wei and Xu 2016; Xu 2013). To reserve a window around the test marker, markers around a ± 2 cM window at the tested marker were excluded from the kinship matrix. The effect of each marker was estimated as a fixed effect and tested using the Wald test statistic (squared effect divided by the variance of the estimated effect). Under the null model, the Wald test follows a chi-square distribution with one degree of freedom, from which a p-value was calculated for each marker. SNP markers of the entire genome were scanned and a test statistic, −log_10_ (*P*-value), profile was plotted. A genome-wide critical value was calculated with a modified Bonferroni correction using a trait specific “effective number of markers or effective degrees of freedom” as the denominator (Wang et al. 2015). The effective degrees of freedom was defined as 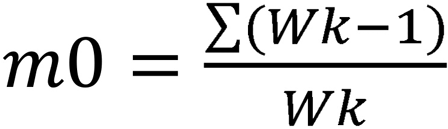, where Wk is the Wald test statistic for SNP k. The trait specific Bonferroni corrected critical value was −log_10_ (0.05/m0). A SNP was declared as significant if its −log_10_ (*P*-value) was larger than −log_10_ (0.05/m0). For each significant SNP, an estimate of the percentage of phenotypic variation was calculated. The proportion of phenotypic variance contributed by each SNP was calculated using the following formula: 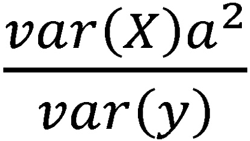, where X is a variable holding the genotype code (+1,−1) for SNP k, var(X) is the variance of variable X, a is the estimated effect for SNP k and var(y) is the total phenotypic variance of the trait under study.

For validation purposes, QTL were also detected by composite interval mapping (CIM; Zeng 1994), using the QTL analysis pipeline of the Breeding Management System (https://www.integratedbreeding.net/).

The physical region of each QTL was determined from the cowpea genome V.1.0 (Lonardi et al. 2017; https://phytozome.jgi.doe.gov/pz/portal.html#!info?alias=Org_Vunguiculata_er), and used to identify candidate genes underlying the QTL interval. Gene models along with their functional annotation were obtained from the Joint Genome Institute cowpea genome portal (https://phytozome.jgi.doe.gov/pz/portal.html#!info?alias=Org_Vunguiculata_er).

## Results

### Phenotypic variation in the population

Nine traits related to domestication in cowpea that differed between the parents were evaluated in the population (Table 1). The means of 100-seed weight, seed number per pod, primary leaf length and width, and pod length of the cultivated parent (IT99K-573-1-1) were all higher than those of the wild parent (TVNu-1158), whereas the mean of peduncle length and days to flowering were higher in the wild parent than the cultivated parent (Table 2). The segregation pattern of pod shattering fit the expected Mendelian ratio of 1:1 (P value = 0.25), while significant segregation distortion was observed for flower color (P value = 0.0035). The frequency of the seven remaining traits followed a normal distribution in the population (Fig.S1). Transgressive segregation was observed for peduncle length, days to flowering, seed number per pod, primary leaf length, and pod length (Table 2).

**Table 2:** Mean, standard error, range, and skewness values for the parental lines and the F8 RIL population.

| Trait | IT99K-573-1-1 | TVNu-1158 | RIL population |  |  |
| --- | --- | --- | --- | --- | --- |
|  |  |  | *Mean +/- SE* | *Range* | *Skewness* |
| Pod shattering | 0 | 1 | - | - | - |
| Peduncle length (cm) | 15.70 +/- 0.98 | 20.73 +/- 1.33 | 16.76 +/- 0.41 | 5.54-33.8 | 0.70 |
| Flower color | 0 | 1 | - | - | - |
| Flowering time (days) | 41.00 ± 1.00 | 70.00 +/- 2.00 | 47 +/- 1 | 35-88 | 1.85 |
| 100-seed weight (g) | 19.6 +/- 0.36 | 0.88 +/- 0.01 | 5.07 +/- 0.17 | 1.77-11.89 | 0.76 |
| Pod length (cm) | 17.89 +/- 1.13 | 6.82 +/- 0.36 | 16.75 +/- 0.40 | 5.07-20.9 | 0.88 |
| Leaf length (cm) | 7.2 +/- 0.33 | 2.76 +/- 0.12 | 4.85 +/- 0.08 | 2.8-8 | 0.23 |
| Leaf width (cm) | 3.94 +/- 0.26 | 1.39 +/- 0.11 | 2.27 +/- 0.05 | 1-4 | 0.65 |
| Number of seeds/pod | 11 +/- 1 | 9 +/- 1 | 9 +/- 1 | 2-16 | 0.06 |

### Development of the cultivated x wild genetic map

A total of 17,739 polymorphic SNPs and 170 RILs were used for genetic map construction. All SNPs were mapped into 1,825 bins on 12 linkage groups (LGs). LGs were named and oriented based on cowpea pseudomolecules (Lonardi et al. 2017), from Vu01 to Vu11. Note that this numbering of LGs differs from the one used in previous cowpea genetic maps (Lucas et al. 2011; Muchero et al. 2009; Muñoz-Amatriaín et al. 2017). A cross reference to the previous nomenclature is included in Table 3. One chromosome (Vu03) was separated into two LGs, which were joined based on the consensus genetic map of Muñoz-Amatriaín et al. (2017). Based on the consensus map, the gap was estimated at 18 cM and was attributed to the absence of polymorphisms between the parents and to the presence of highly distorted markers. The genetic map covered 1,026.03 cM, with an average density of one marker bin per 1.8 cM, and 9.72 SNPs per bin. The length of the LGs varied from 15.25 (Vu04) to 139.72 cM (Vu03) (Table 3). This genetic map contains the highest number of SNPs in an individual cowpea map to date. Compared to the previously published genetic maps of cowpea constructed with SNP markers (Lucas et al. 2011; Muchero et al. 2009; Muñoz-Amatriaín et al. 2017), we observed a much shorter Vu04 (old LG11). The difference may be due to genomic divergences between the cultivated and wild parents leading to disturbed chromosome pairing which reduces or suppresses recombination in affected regions and/or structural chromosomal rearrangement. The IT99K-573-1-1 x TVNu-1158 genetic map can be found in supplementary information (Table S1).

**Table 3:** Number of SNPs, and length of each linkage group in the cultivated x wild cowpea genetic map.

| **Linkage Group (LG, old)** | **Chromosome** | **SNPs** | **Bins** | **cM** |
| --- | --- | --- | --- | --- |
| 4 | Vu01 | 1742 | 132 | 65.31 |
| 7 | Vu02 | 1368 | 140 | 98.62 |
| 3 | Vu03 | 2780 | 280 | 157.61 |
| 11 | Vu04 | 117 | 24 | 15.25 |
| 1 | Vu05 | 1740 | 205 | 114.62 |
| 6 | Vu06 | 1676 | 150 | 78.29 |
| 2 | Vu07 | 1968 | 228 | 134.15 |
| 5 | Vu08 | 1607 | 159 | 81.59 |
| 8 | Vu09 | 1774 | 189 | 100.37 |
| 10 | Vu10 | 1416 | 144 | 77.37 |
| 9 | Vu11 | 1551 | 174 | 102.85 |
| **Total** | **Total** | **17739** | **1825** | **1026.03** |

### Identification of domestication-related QTL and candidate genes

Two mapping methods were used for QTL analysis (see Materials and Methods for details). CIM detected a total of 27 QTL for all traits, while the mixed model for QTL mapping (Xu 2013) identified 22 QTL, all of which were also identified by CIM. A few of these 22 QTL were no longer significant after Bonferroni correction (α =0.05). In this paper, only 16 significant QTL from the mixed model are reported and discussed further. The 16 domestication-related QTL were distributed on seven of the eleven chromosomes of cowpea (Table 4; Fig. 1) and their – log_10_ (*P*-value) ranged from 5.15 for seed number per pod to 20.00 for peduncle length, flower color, and leaf width (Table 4). The significance levels ranged from 4.57 to 4.98 depending on the trait (Table S2). The percentage of phenotypic variation for the identified QTL ranged from 18.32% for seed number per pod to 85.65% for flower color (Table 4). We identified four regions showing QTL clustering for domestication traits, including one region on Vu08 where four QTL related to increased organ size (seed weight, pod length, leaf length and leaf width) were mapped. Further studies would be required to determine if this clustering of domestication-related QTL results from pleiotropic effects or tightly linked QTL.

**Table 4:** QTL for the domestication-related traits identified by the linear mixed model analysis and their map position.

| Trait | QTL<sup>a</sup> | Peak SNP | Chr V u | Position (cM) | Consensus map position (cM) | - Log10(P) | QTL region (cM) | % Phenotypic variation | Effect<sup>b</sup> |
| --- | --- | --- | --- | --- | --- | --- | --- | --- | --- |
| Pod shattering | *CPsh at3* | 2_45094 | 03 | 104.84 | 94.54 | 8.17 | 100.57-106.67 | 37.69 | -0.31 |
|  | *CPsh at5* | 2_23044 | 05 | 100.02 | 73.45 | 7.28 | 96.11 - 103.85 | 30.27 | -0.28 |
| Peduncle length | *CPed l5* | 2_00251 | 05 | 66.23 | 57.10 | 20 | 60.97-76.34 | 71.83 | -4.50 |
| Flower color | *CFco l7* | 2_24259 | 07 | 65.89 | 31.60 | 20 | 64.39-72.43 | 85.65 | -4.16 |
| Flowering time | *CFt5* | 2_05332 | 05 | 5.77 | 3.20 | 5.51 | 0.93-7.93 | 20.00 | 4.13 |
|  | *CFt9* | 2_03945 | 09 | 21.79 | 22.15 | 20 | 13.82-30.02 | 79.30 | -8.46 |
| 100-seeds weight | *CSw1* | 2_17042 | 01 | 46.62 | - | 6.85 | 45.15-49.45 | 19.85 | 1.03 |
|  | *CSw6* | 2_55420 | 06 | 72.16 | - | 6.66 | 69.26-73.30 | 21.48 | 1.06 |
|  | *CSw8* | 2_05809 | 08 | 78.57 | 72.39 | 11.86 | 76.14-81.59 | 36.87 | 1.39 |
| Pod length | *CPod l3* | 2_49508 | 03 | 116.49 | 101.37 | 5.19 | 116.02-116.49 | 23.98 | 1.08 |
|  | *CPod l8* | 2_00195 | 08 | 78.17 | - | 13.04 | 74.67-81.59 | 46.08 | 1.52 |
| Leaf length | *CLl8* | 2_09764 | 08 | 78.87 | - | 6.98 | 78.17-80.02 | 38.24 | 0.69 |
| Leaf width | *CLw1* | 2_19941 | 01 | 34.13 | 38.65 | 20 | 30.35-49.45 | 63.28 | 0.54 |
|  | *CLw8* | 2_21200 | 08 | 78.87 | - | 7.09 | 76.73-79.18 | 34.75 | 0.40 |
| N° seed per pod | *CSp5* | 2_07573 | 05 | 11.37 | 6.79 | 5.15 | 10.57-12.11 | 18.32 | 1.07 |
|  | *CSp9* | 2_19332 | 09 | 24.73 | - | 5.77 | 22.51-25.32 | 21.09 | -1.18 |

<sup>a</sup> QTL name is designated as follow: “C” to indicate cowpea, followed by the trait code, then followed by the chromosome number
<sup>b</sup> Positive or negative effect alleles, for which a positive value indicates allele of the cultivated parent is positive, and a negative value indicates the allele of the wild parent is positive.

**Fig. 1:**
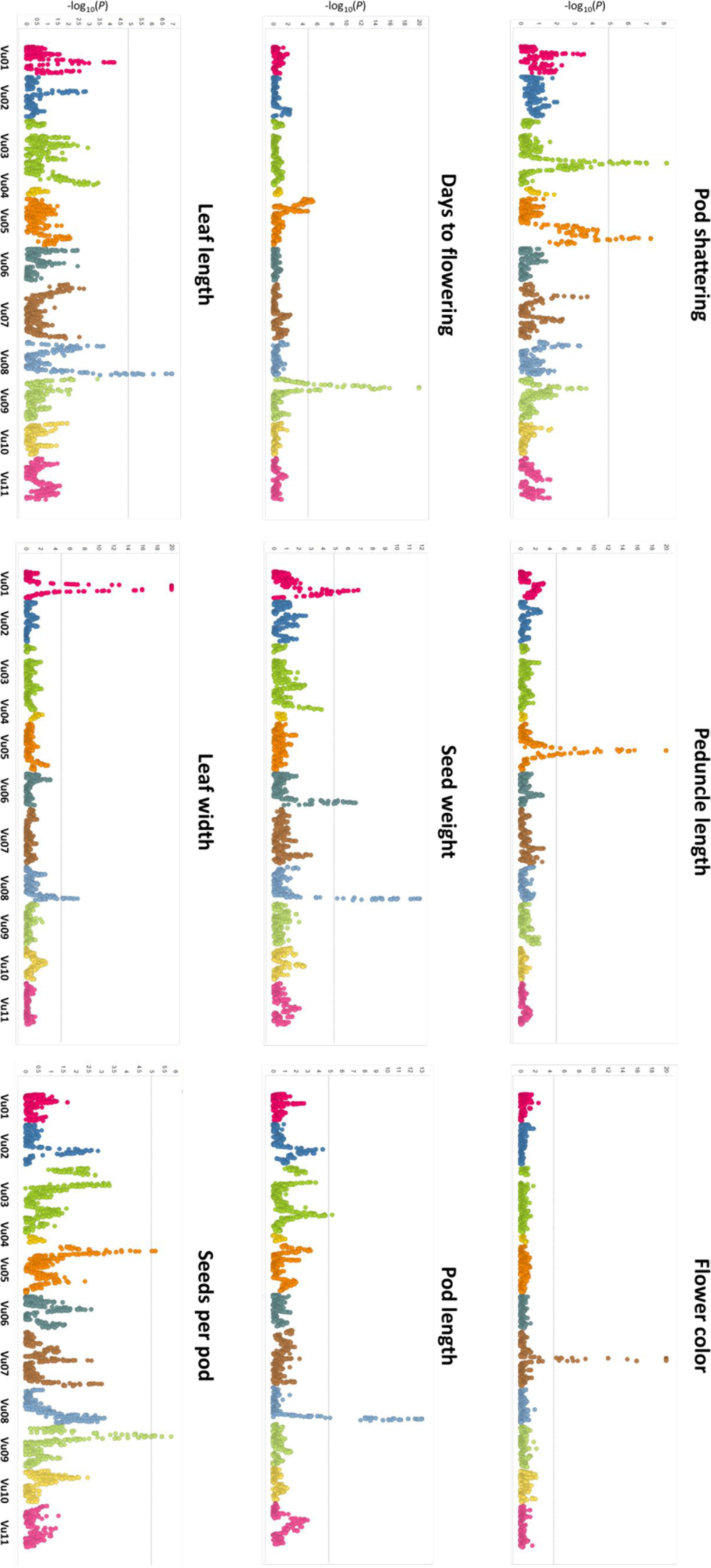
Genetic map of the RILs and QTL plots for the nine domestication-related traits. The horizontal axis indicate the chromosomes, the vertical axis indicates the −log10 of the probability (*P*-values). The dashed line indicates the significance threshold at 0.05.

*Pod shattering.* Two significant QTL were detected, on Vu03 and Vu05, for pod shattering (Table 4; Fig. 1). These QTL named *CPshat3* and *CPshat5* explained 37.69 % and 30.27 % of the phenotypic variation, respectively. *CPshat3* spanned 6.1 cM corresponding to ~ 3.4 Mb on the cowpea pseudomolecules (Lonardi et al. 2017) and contained 268 genes, while *CPshat5* spanned 7.74 cM corresponding to ~ 1.60 Mb and contained 204 annotated genes (Table S3). Among the genes underlying the main QTL (*CPshat3*) there were two with a possible role in pod shattering: *Vigun03g306000*, which encodes a NAC-domain transcription factor (NAC007) involved in secondary cell wall biosynthesis (Wang et al. 2011), and *Vigun03g302600*, encoding a C2H2-like zinc finger protein. Members of this family have been found to regulate lignin accumulation in *Brassica napus* (Tao et al. 2017). Among the annotated genes underlying the *CPshat5* region, *Vigun05g273500*, annotated as Myb domain protein 26, is the best candidate. An orthologue of *Vigun05g273500* in Arabidopsis has been associated with anther dehiscence (Yang et al. 2007).

*Peduncle length*. QTL analysis detected one major QTL on Vu05, *CPedl5*. This QTL accounted for 71.83% of the phenotypic variation and it spanned a 15.37 cM region corresponding to ~ 4.53 Mb (Table 4; Fig.1). A total of 379 annotated genes were identified in the *CPedl5* interval (Table S3). Notably, three genes encode auxin-responsive GH3 family proteins (*Vigun05g201700*, *Vigun05g217600* and *Vigun05g223100*) and members of the GH3 family have been found to influence organ elongation (Nakazawa et al. 2001; Takase et al. 2004).

*Flower pigmentation.* A major-effect QTL controlling purple flower (*CFcol7*) was detected in a 64 cM region (~ 4.56 Mb) on Vu07 (Table 4; Fig.1). This QTL explained 85.65% of the phenotypic variation and contained 254 annotated genes (Table S3). The transcription factor TRANSPARENT TESTA8 (TT8, *Vigun07g110700*), which contains a basic helix-loop-helix domain, was identified in the QTL region. TT8 is known to be involved in the regulation of flavonoid biosynthesis in Arabidopsis (Nessi et al. 2000).

*Time to flowering.* Two significant QTL related to time to flowering were detected, on Vu05 and Vu09. The QTL on Vu05 (*CFt5*) explained 20% of the phenotypic variation, while *CFt9* on Vu09 explained 79.3% of the phenotypic variation (Table 4; Fig.1). *CFt5* spans 7 cM which correspond to ~ 6.64 Kb, while *CFt9* maps to a 16 cM region corresponding to ~ 3.86 Mb. A total of 86 genes were identified in the *CFt5* region, while 299 genes were identified underlying *CFt9* (Table S3). Among the annotated genes in the *CFt9* region, a phytochrome E photoreceptor (*Vigun09g050600*), and a transcription factor TCP 18 (*Vigun09g062200*) were found. TCP 18 has been shown to delay flowering by repressing differentiation of axillary buds (Niwa et al. 2013). No genes with obvious roles in flowering were identified in the *CFt5* region (Table S3).

*100-seed weight.* Three QTL for seed weight were detected on Vu01, Vu06, and Vu08. The QTL with the highest contribution to the trait (*CSw8*) was located on Vu08 and explained 36.87% of the phenotypic variation, while *CSw1* and *CSw6* explained 19.85% and 21.48% of the phenotypic variation, respectively (Table 4). *Csw8* spans 5.45 cM (~ 1.61 Mb) and contains 225 genes, while *Csw1* and *Csw6* span 4.3 cM (~1.60 Mb) and 4.04 cM (~1.12 Mb) and contain 206 and 160 annotated genes, respectively (Table S3). Among the many genes in those QTL region, several genes involved in carbohydrate metabolism including UDP-glycosyltransferases and cellulose synthases were identified.

*Pod length*. Pod length was analyzed as a measure of increase in organ size, and two QTL were identified on Vu03 and Vu08. These QTL explained 23.98% and 46.08% of the phenotypic variation, and spanned 0.47 cM (~ 84 Kb) and 6.92 cM (~ 2.1 Mb), respectively. Nine genes including an AGAMOUS-like8 (*Vigun03g343500*) which is known to mediate cell differentiation during fruit development in Arabidopsis (Gu et al. 1998) were identified underlying *CPodl3*, while 309 genes were found in the *CPodl8* interval. *CPodl8* maps to the same genomic region as *CSw8* (Table 4; Fig. 1) and includes genes encoding cellulose synthases, UDP-Glycosyltransferases and a cluster of pectin lyases (Table S3), all involved in carbohydrate metabolism.

*Leaf length and width.* The leaf is the main organ for photosynthesis in cowpea and one QTL for leaf length (*CLl8*) and two QTL for leaf width (*CLw1*, *CLw8*) were identified in this population (Table 4; Fig. 1). *CLl8* explained 38.24% of the phenotypic variation and spanned 1.85 cM (~ 9.9 Kb). *CLw1* accounted for 63.28% of the phenotypic variation and spanned 19.1 cM (~ 3.61 Mb), while *CLw8* explained 34.75% of the phenotypic variation and spanned 2.45 cM (~1.08 Mb). Regarding leaf length, a total of 142 genes were identified in the physical region of *CLl8.* For leaf width, over 300 genes were identified in *CLw1* interval, while 148 genes were identified underlying *CLw8* (Table S3). *CLl8* and *CLw8* map to the same genomic region as *CSw8* and *CPodl8* (Table 4; Fig. 1). Genes present in *Csw8* and *CPodl8* are also contained in these two QTL, including proteins of the pectin lyase-like superfamily and cellulose synthases (Table S3).

*Seed number per pod.* Two QTL were detected, on Vu05 and Vu09, accounting for 18.32% and 21.09% of the phenotypic variation, respectively and spanned 1.54 cM (~3.71 Kb) and 2.81 cM (~4.94 Kb)respectively. The total number of genes underlying *CSp5* and *CSp9* was only 40 and 33, respectively. One of the genes underlying *CSp9* encodes a transcription factor TCP5, which has been associated with ovule development in Arabidopsis (Wei et al. 2014). We did not identify any gene in the *CSp5* region with known functions related to seed number per pod (Table S3).

## Discussion

Through the domestication process, cowpea underwent many phenotypic changes compared to its wild progenitor. In this study, we have focused on nine traits that differ between wild and cultivated cowpea and identified 16 QTL for trait determination distributed over seven chromosomes. For each trait, only one or two major QTL were identified except for seed weight, for which three QTL were detected. These results are consistent with previous studies suggesting that, in most crops, domestication related traits seem to be controlled by a small number of QTL with large phenotypic effects (reviewed by Burger et al. 2008, and Gross and Olsen 2010).

### Identification of novel QTL for pod shattering and time to flowering

Among the domestication traits considered in this study, loss of pod shattering and time to flowering are the most relevant for cowpea breeding, together with increase of organ size. The pod shattering habit causes pre-harvest yield losses. Two major QTL were identified for pod shattering, on Vu03 and Vu05. In previous studies in cowpea, Suanum et al. (2016) reported one major QTL and one minor QTL for pod shattering, while Andargie et al. (2014) identified only one QTL explaining 10.8% of the phenotypic variation. While it was not possible to compare the QTL identified by Suanum et al. (2016) because of unavailability of marker sequences, BLAST searches of SSR markers from the Andargie et al. (2014) study to the reference genome sequence (https://phytozome.jgi.doe.gov/pz/portal.html#!info?alias=Org_Vunguiculata_er)revealed that the QTL identified in that study is located in a different chromosome compared to those reported here. This suggests the QTL identified in this study are novel.

Genes involved in seed dispersal have been cloned in dicots including soybean (Dong et al. 2014), Brassica (Tao et al. 2017) and Arabidopsis (Liljegren et al. 2000). For *CPshat3*, the major pod shattering QTL, we identified a gene (*Vigun03g306000*) encoding a NAC domain transcription factor. In soybean, the NAC family gene *SHATTERING1-5* has been found to activate secondary wall biosynthesis affecting pod shattering resistance (Dong 2014). In addition, a C2H2 zinc finger protein (*Vigun03g302600*) was identified in this QTL region. A member of the C2H2 zinc finger family of proteins has been shown to enhance silique shattering resistance in Brassica (Tao et al. 2017). In the *CPshat5* region we identified *Vigun05g273500*, a gene annotated as Myb domain protein 26. *Vigun05g273500* is an ortholog of *AT3G13890.1*, which acts upstream of the lignin biosynthesis pathway and has been shown to be key for anther dehiscence by regulating the development of secondary thickening in the endothecium in Arabidopsis (Yang et al. 2007).

Time to flowering is one of the most important agronomic traits that plays a key role in the adaptation of a variety to specific agro-ecological areas. Two major QTL associated with time to flowering have been identified in this work. Similarly to pod shattering, none of these QTL seem to coincide with the main QTL reported by Andargie et al. (2014). The main QTL for time to flowering (*CFt9*) in the present study mapped to Vu09, with the wild accession alleles conferring late flowering. There are two genes in the *CFt9* region with functions related to time to flowering. One is a phytochrome E (*Vigun09g050600*), one of the photoreceptors perciving red/far-red light ratio and influencing flowering time (Devlin et al. 1998). *Vigun09g050600* is an ortholog of the adzuki bean *Vigan.02G285600.01*, which is one of the candidate genes for the major photoperiod QTL *Flowering Date1* (Yamamoto et al. 2016). The other gene is *Vigun09g062200*, encoding a transcription factor TCP 18 which has been previously shown in Arabidopsis to repress flowering by interacting with the florigen proteins FLOWRING LOCUS T (FT) and TWIN SISTER OF FT (TSF) (Niwa et al. 2013).

### QTL clusters controlling increased organ size

Larger seeds play a major role in consumer preference, and larger leaves provide more surface than smaller leaves for the production of photosynthate. Four traits including 100-seed weight, pod length, primary leaf length and width were analyzed as a measure of increased organ size in the population. QTL identified for seed weight (*CSw8*), leaf length (*CLl8*), leaf width (*CLw8*), and pod length (*CPodl8*) mapped to the same region on Vu08. Multiple QTL located in this region suggest potential pleiotropy or clustering of genes controlling increased organ size, a fundamental target during domestication. In rice bean, QTL for leaf size were detected close to QTL controlling seed- and pod-size related traits (Isemura et al. 2010). A cluster of pectin lyase-like superfamily proteins, which are known to be involved in many biological processes including cellular metabolism (Cao 2012) was found in the QTL region for 100-seed weight, pod length, and leaf size. Also, clusters of other genes with functions in carbohydrate metabolism such as UDP-glycosyltransferases and cellulose synthases were identified in this region. Further work, including fine mapping of this hotspot region, would be needed to elucidate the genetic control of increased organ size in cowpea.

Two additional QTL related to seed weight were also detected outside this region, on Vu01 and Vu06. One of them (*CSw6*) mapped to the same region as *Css-4*, a seed size QTL identified from a cultivated by cultivated RIL population (Lucas et al. 2013). There are two other previous studies on cowpea reporting QTL involved in seed weight using wild x cultivated population (Andargie et al. 2014; Fatokun et al. 1992). While it was not possible to compare QTL regions between this study and that of Fatokun et al. 1992 because of unavailability of marker sequences, none of these QTL seem to coincide with the ones reported by Andargie et al. 2014.

### Genetic control of other domestication traits

As one of the major yield components, the number of seeds per pod, is an important trait in cowpea breeding. Two QTL, *CSp5* and *CSp9*, were detected for this trait on Vu05 and Vu09, respectively. The main QTL (*CSp9*) accounted for 21.09% of the phenotypic variation and co-localized with the main QTL for days to flowering. The allele from the wild parent at *CSp9* conferred a higher number of seeds per pod, then making it a good target for introgression into cultivated cowpea varieties. *Vigun09g060700* was identified in *Csp9* region. This gene is annotated as a transcription factor TCP5, which is involved in ovule development in Arabidopsis (Wei et al. 2014). Since the number of seeds is determined among other things by the number of ovules (and fertile ovules) per ovary, this gene is a promising candidate for number of seeds per pod in cowpea.

Another major difference observed between the wild and the cultivated parent used in this study was the length of the peduncles. A single QTL with a high contribution to the phenotypic variation of the trait (71.83%) was identified on Vu05. Long peduncles in cowpea are desirable as they allow pods to be positioned above the canopy, a characteristic that reduces damage to pods by the pod borer *Maruca vitrata* and are also advantageous for harvesting of pods. Some genes involved in plant growth and development were found underlying *CPedl5*, including genes belonging to the auxin-responsive GH3 family. In cucumber, two genes belonging to this family (*Csa6G492310* and *Csa6G493310*) were identified as candidates for the main QTL for fruit peduncle length (Song et al. 2016).

The presence or absence of purple pigment in the flower is controlled by a single major QTL in this population. This is similar to a previous study in soybean, where one QTL for flower color was identified (Josie et al. 2007). The QTL controlling flower color was mapped to Vu07 and explained 85.65% of the phenotypic variation for the trait. An earlier study of cowpea suggested that a single gene controls flower color with purple flower color being dominant (Sangwan and Lodhi 1998). *Vigun07g110700*, involved in flavonoid biosynthesis, was identified in the Q TL region. *Vigun07g110700* has sequence similarity to the Arabidopsis *TT8* gene (*AT4G09820.1*) encoding a basic helix-loop-helix domain protein involved in the control of flavonoid biosynthesis, whose final compounds include anthocyanins (Nesi et al. 2000). *Medicago truncatula TT8 (MtTT8)* has also been found to regulate anthocyanin and proanthocyanidin biosynthesis by interacting with other transcription factors and forming regulatory complexes (Li et al. 2016). Hence, *Vigun07g110700* is a strong candidate for the control of flower pigmentation in cowpea.

In summary, this study provides novel QTL for many DRTs as well as a link between genetic and physical maps. Thanks to the new genomic resources available for cowpea, which include a reference genome sequence of IT97-499-35 (Lonardi et al. 2017; available from Phytozome), we were able to estimate physical sizes for all QTL identified and to determine, for the first time in cowpea, candidate genes underlying QTL controlling DRTs. The results of this study provide a basis for further fine mapping of genes involved in cowpea domestication and a genetic foundation for the utilization and exploitation of wild relatives in cowpea breeding programs.

## Author contribution statement

CF, OB, NC, TC and PR designed the experiments and edited the manuscript. CF, and OB performed the phenotypic data collection. SL, and YG extracted the DNA from the RIL population for genotyping. SL conducted data analysis, and wrote the paper. MMA, IH, and SX participated in data analysis. CF, SX and MMA assisted in writing the paper. All authors read and approved the final manuscript.

## Acknowledgments

This work was supported by grants from the Generation Challenge Program (TL1), and Feed the Future Innovation Lab for Climate Resilient Cowpea (Cooperative Agreement AID-OAA-A-13-00070). Sassoum Lo was supported by funds from the West Africa Agricultural Productivity Program. We thank the International Institute of Tropical Agriculture for the RIL population. We also thank Abdou Souleymane (INRA, Niger) for helping to identify the wild parent, Stefano Lonardi and Steve Wanamaker (University of California Riverside, USA) for the cowpea genome sequence and annotations, and Dr. Paul Gepts (University of California Davis, USA) for his valuable inputs.

### Conflict of interest

The authors declare that they have no conflict of interest.

---

**Fig. S1:**
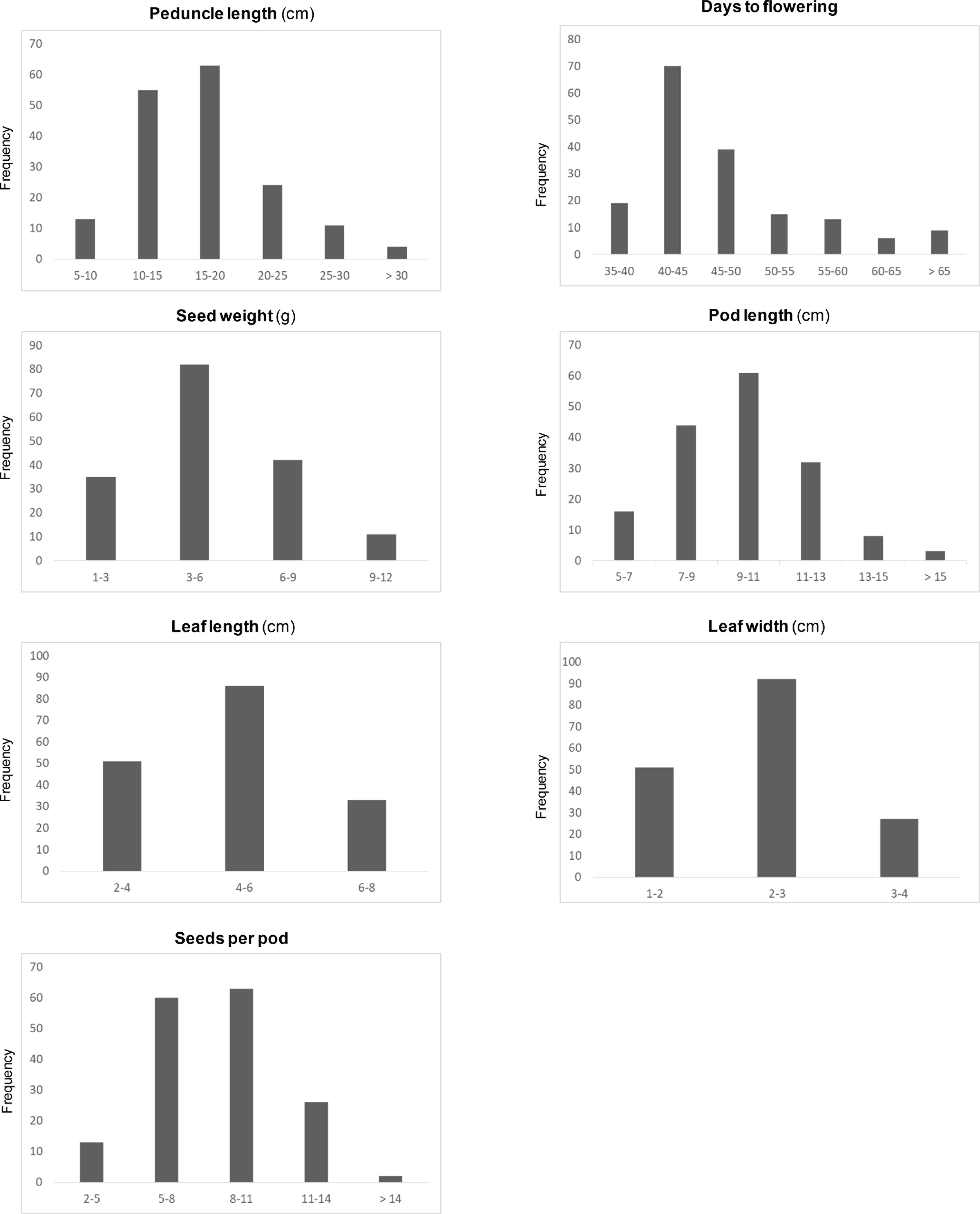
Phenotypic distribution of seven out of the nine domestication-related traits evaluated in the F8 population.

